# Rapid CD4 cell loss is caused by specific CRF01_AE cluster with V3 signatures favoring CXCR4 usage

**DOI:** 10.1101/427625

**Authors:** Hongshuo Song, Weidong Ou, Yi Feng, Junli Zhang, Fan Li, Jing Hu, Hong Peng, Hui Xing, Liying Ma, Qiuxiang Tan, Beili Wu, Yiming Shao

**Affiliations:** State Key Laboratory for Infectious Disease Prevention and Control, National Center for AIDS/STD Control and Prevention, Chinese Center for Disease Control and Prevention, Beijing 102206, China; Chinese Academy of Sciences, Shanghai 201203, China; Center of Infectious Diseases, Peking University, Beijing 100191, China

**Keywords:** CRF01_AE, Coreceptor tropism, CXCR4, HIV-1 pathogenesis

## Abstract

HIV-1 evolved into various genetic subtypes and circulating recombinant forms (CRFs) in the global epidemic, with the same subtype or CRF usually having similar phenotype. Being one of the world’s major CRFs, CRF01_AE infection was reported to associate with higher prevalence of CXCR4 (X4) viruses and faster CD4 decline. However, the underlying mechanisms remain unclear. We identified eight phylogenetic clusters of CRF01_AE in China and hypothesized that they may have different phenotypes. In the national HIV molecular epidemiology survey, we discovered that people infected by CRF01_AE cluster 4 had significantly lower CD4 count (391 vs. 470, *p* < 0.0001) and higher prevalence of predicted X4-using viruses (17.1% vs. 4.4%, *p* < 0.0001) compared to those infected by cluster 5. In a MSM cohort, X4-using viruses were only isolated from sero-convertors infected by cluster 4, which associated with rapid CD4 loss within the first year of infection (141 vs. 440, *p* = 0.01). Using co-receptor binding model, we identified unique V3 signatures in cluster 4 that favor CXCR4 usage. We demonstrate for the first time that HIV-1 phenotype and pathogenicity can be determined at the phylogenetic cluster level in a single subtype. Since its initial spread to human from chimpanzee in 1930s, HIV-1 remains undergoing rapid evolution in larger and more diverse population. The divergent phenotype evolution of two major CRF01_AE clusters highlights the importance in monitoring the genetic evolution and phenotypic shift of HIV-1 to provide early warning for the appearance of more pathogenic strains such as CRF01_AE cluster 4.

**Significance Statement:** Past studies on HIV-1 evolution were mainly at the genetic level. This study provides well-matched genotype and phenotype data and demonstrates disparate pathogenicity of two major CRF01_AE clusters. While both CRF01_AE cluster 4 and cluster 5 are mainly spread through the MSM route, cluster 4 but not cluster 5 causes fast CD4 loss, which is associated with the higher prevalence CXCR4 viruses in cluster 4. The higher CXCR4 use tendency in cluster 4 is derived from its unique V3 loop favoring CXCR4 binding. This study for the first time demonstrates disparate HIV-1 phenotype between different phylogenetic clusters. It is important to monitor HIV-1 evolution at both the genotype and phenotype level to identify and control more pathogenic HIV-1 strains.

## Introduction

During global transmission, HIV-1 evolved into various subtypes and their hybrids, the so called circulating recombinant forms (CRFs) (1). CRF01_AE, one of the major CRFs spreads mainly in Southeast Asia and China (1, 2). Early studies in Thailand reported faster CD4 loss and shorter survival of CRF01_AE infected people compared to infections in western countries where subtype B predominate (3–6). In recent years, faster disease progression of CRF01_AE was also reported in China (7–9). Although past studies reported high prevalence of X4 viruses and fast disease progression of CRF01_AE infections (7, 8, 10, 11), the data were mainly based on samples without known infection time and genotypic prediction without phenotypic confirmation. It is still unclear how early X4 viruses can emerge during natural CRF01_AE infection and whether the genotypic prediction is reliable.

The epidemic of CRF01_AE in China was initiated by multiple phylogenetic clusters introduced from Thailand in 1990s (2). We formerly identified eight CRF01_AE clusters, which have different geographic distributions and epidemic patterns (2, 12). Therefore, we hypothesized that they may have different phenotypes. We used the national HIV molecular epidemiology survey (NHMES) dataset for screening and a men-who-have-sex-with-men (MSM) sero-incidence cohort for in-depth study to determine the time of X4 virus emergence, and phenotypically confirmed viral tropism using matched viral isolates. We observed significantly lower CD4 counts in CRF01_AE cluster 4 compared to cluster 5 in the NHMES dataset. Focusing on the MSM cohort, we demonstrated higher prevalence of X4 phenotype in cluster 4 among people within the first year of infection. We further determined the genetic and structural basis favoring X4 co-receptor usage in cluster 4. This is the first demonstration that different clusters from the same HIV-1 subtype can cause disparate rate of disease progression and bridged the missing link between CRF01_AE genetic makeup, phenotype and clinical outcomes.

## Results

### Lower CD4 T cell count and higher prevalence of X4-using virus in CRF01_AE cluster 4

We compared the CD4 T cell counts of 1118 CRF01_AE, 633 CRF07_BC and 123 subtype B newly diagnosed HIV-1 positive participants in the China’s NHMES study and found no differences (Fig. 1A). When analyzing the two major CRF01_AE clusters which contribute greatly to China’s MSM epidemic, however, the CD4 count in cluster 4 was significantly lower than in cluster 5 (p < 0.0001) (Fig. 1B). Since CD4 counts decline with time during natural infection, we distinguished cases of recent infection from long-term infection by HIV-1 Limiting Antigen-Avidity assay. Significantly lower CD4 count in cluster 4 was found in both recent and long-term infection groups (*p* = 0.0004 and *p* < 0.0001, respectively) (Fig. 1C-D). The proportion of people with CD4 below 200 was higher in cluster 4 than in cluster 5 (9.2 % vs.1.8 %, *p* = 0.0001) (Fig. 1B), while the rates of recent infection were nearly the same between the two clusters (31.3% vs. 31.4%). The results clearly showed that the CRF01_AE cluster 4 but not cluster 5 leads to rapid CD4 cell loss after infection.

**Figure 1.**
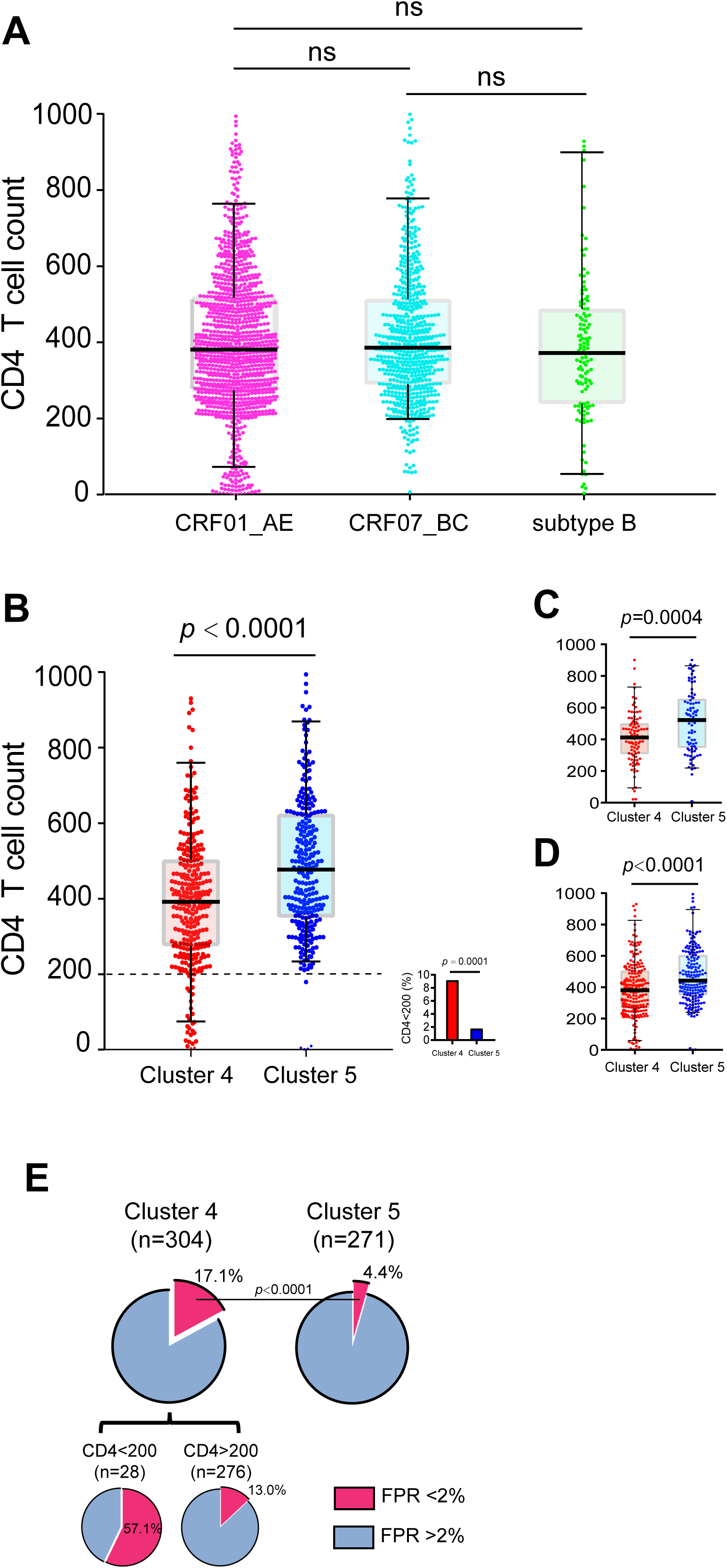
Comparison of CD4 T cell count and prevalence of X4 virus in different HIV-1 subtypes and CRF01_AE clusters. (A) Comparison of CD4 T cell count between individuals infected by CRF01_AE (n=1118), CRF07_BC (n=633) and subtype B (n=123) from the national HIV molecular epidemiology survey. (B-D) Significantly lower CD4 T cell count among individuals infected by CRF01_AE cluster 4 (n=308) than those infected by cluster 5 (n=273) regardless of the stage of infection (B), in the recent infection group (C), and in long-term infection group (D). The small figure in panel B shows the percentage of individuals with CD4 below 200. In each panel, the vertical line, box and whisker represents the median, upper and lower quantiles, and the 5–95 percentile, respectively. The statistical difference in CD4 count between different groups was calculated using two-tailed Mann-Whitney U test. The percentage of subjects with CD4 below 200 was compared using two-tailed Fisher’s exact test. (E) Prevalence of predicted X4 viruses in CRF01_AE cluster 4 and cluster 5, and in the lower (<200) and higher (>200) CD4 groups in CRF01_AE cluster 4. The statistical difference was determined using two-tailed Fisher’s exact test.

Because CXCR4 tropism is associated with lower CD4 cell count (13–17), we investigated the prevalence level of X4 viruses in clusters 4 and 5 using genotypic prediction. Geno2pheno prediction (FPR cutoff = 2%) showed higher prevalence of X4 genotype in cluster 4 than in cluster 5 (17.1% vs. 4.4%, *p* < 0.0001) (Fig. 1E). In cluster 4, the X4 genotype were "concentrated” among patients with low CD4 counts (Fig. 1E). This result indicates a strong association between higher X4 prevalence and lower CD4 count in CRF01_AE cluster 4 infection.

### High prevalence of X4-using phenotype in CRF01_AE cluster 4 among recently infected individuals

To further characterize the mechanism of fast CD4 cell loss in CRF01_AE cluster 4, we focused on a sero-incidence cohort with around two thousands of MSMs from Beijing Chaoyang District (the CYM cohort). For the 135 MSM sero-convertors, we first performed Illumina deep sequencing on all 78 participants with archived first year blood samples and successfully sequenced 71 of them, with an average of 65,000 sequencing reads per participant (SI Appendix, Figs. S1-S2, Tables S1-S2). Analysis of Geno2pheno FPR distribution among the plasma viral quasispecies in each participant again found higher frequency of X4-using variants in people infected by CRF01_AE cluster 4 than by cluster 5, CRF07_BC and subtype B (Fig. 2 and SI Appendix, Fig. S3 and Table S2). Together, deep sequencing on samples collected within the first year of infection confirmed the observation in large cross-sectional NHMES study.

**Figure 2.**
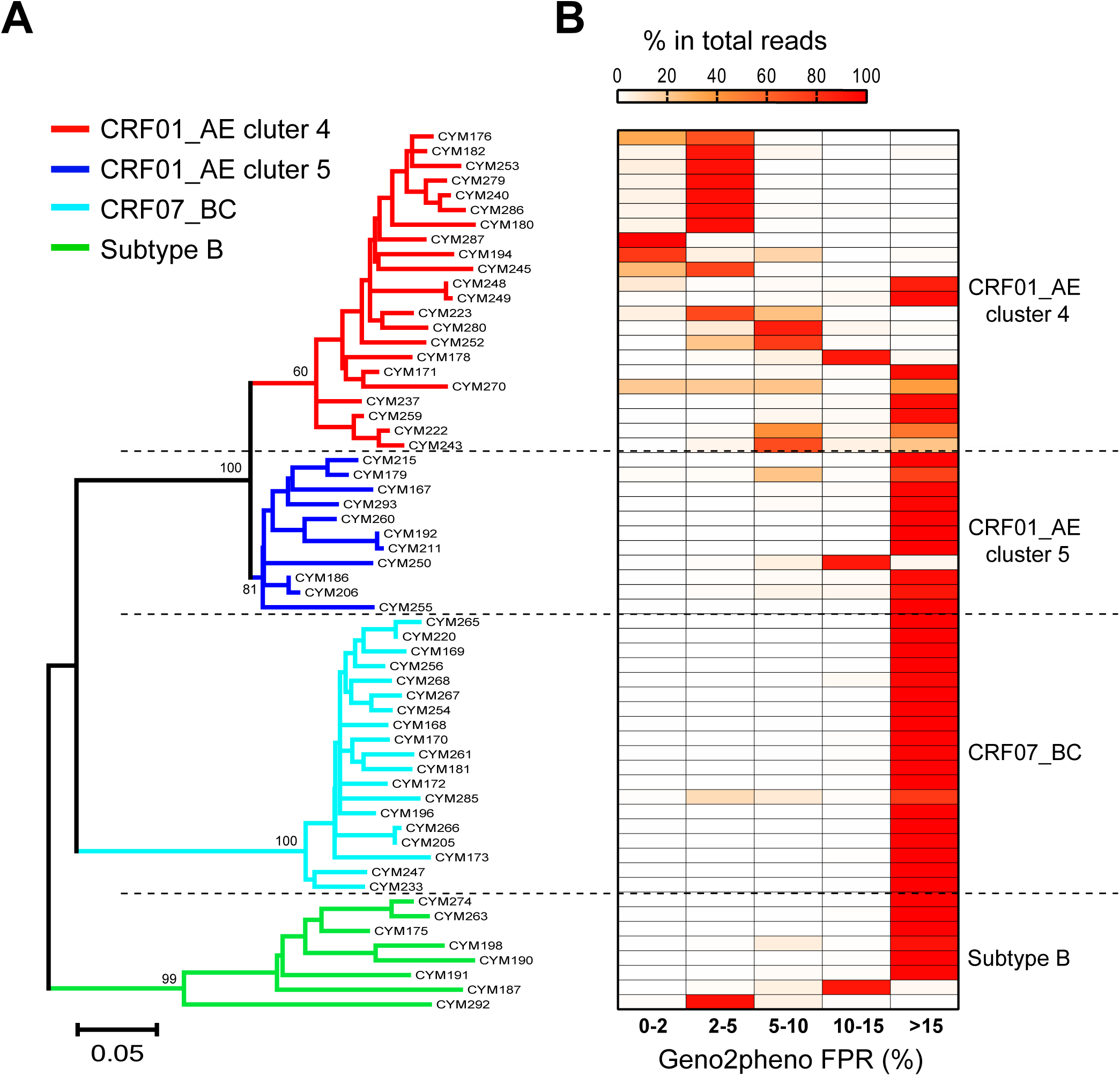
Higher frequency of predicted X4-using variants in CRF01_AE cluster 4 identified by deep sequencing. (A) Phylogenetic relationship of 60 deep sequenced individuals from the CYM cohort. In each individual, the most frequent haplotype among the deep sequencing reads was used for phylogenetic inference. The Neighbor-joining (NJ) tree was constructed using the Kimura 2-parameter evolutionary model with 1000 bootstrap replications. In the tree, the branches for CRF01_AE cluster 4 (n=22), cluster 5 (n=11), CRF07_BC (n=19) and subtype B (n=8) were color coded. (B) Heatmap showing the frequency distribution of Geno2pheno FPR value among the deep sequencing reads in each individual. The samples in the phylogenetic tree and in the heatmap were matched.

To confirm the genetic prediction phenotypically, we isolated viruses using cryopreserved PBMC from the deep sequenced CRF01_AE participants and conducted coreceptor tropism assays using GHOST cell lines. A total of 24 viruses were successfully isolated. Five showed X4-using phenotype and the remaining 19 were R5-only phenotype (Fig. 3A-C). All of the five isolates with X4-using phenotype, including four dual-tropic and one exclusively X4-tropic belonged to cluster 4 (Fig. 3A and SI Appendix, Fig. S4) (The method to determine coreceptor tropism was described in Materials and Methods and Fig. S4). This indicates a much higher prevalence of X4-using phenotype in cluster 4 than in cluster 5 (31.3% vs. 0%), consistent with the result of genotypic prediction.

**Figure 3.**
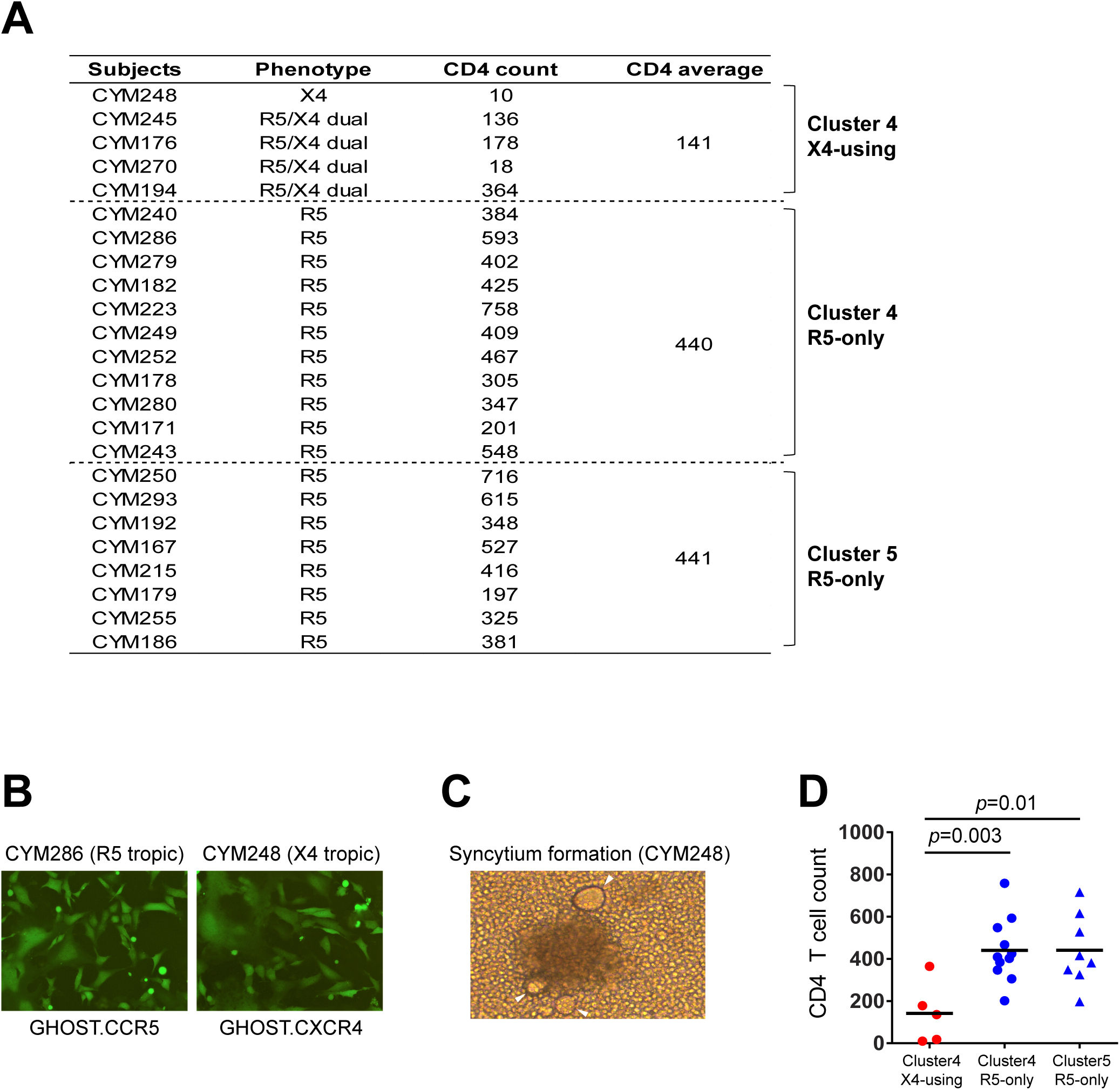
Phenotypic characterization of primary CRF01_AE viral isolates and the association between coreceptor tropism and CD4 count. (A) Coreceptor usage phenotype of 24 primary viral isolates from CRF01_AE cluster 4 (n=16) and cluster 5 (n=8) and the corresponding CD4 counts in each individual. (B) Determination of coreceptor tropism using GHOST.CCR5 and GHOST.CXCR4 cell lines. The GFP expression induced by one representative R5-only isolate and one X4-using isolate were shown. (C) Syncytium formation at day 12 post PBMC co-cultivation in the culture of CYM248, an isolate using CXCR4 exclusively. (D) Significantly lower CD4 count in individuals harboring X4-using viruses in CRF01_AE cluster 4. The black vertical line represents the median CD4 count. The statistical difference was calculated using two-tailed Mann-Whitney U test.

### Individuals harboring X4 viruses had significantly lower CD4 T cell count

Previous studies demonstrated the association between X4-using phenotype and lower CD4 count, mainly for subtype B HIV-1 (13–17). However, a recent study based solely on genotypic prediction failed to find such an association in CRF01_AE infected people in China (though there is a trend that people with CD4<50 tend to have Geno2pheno FPR<5) (8). Because genotypic prediction could overestimate the actual X4 prevalence, we therefore compared the CD4 T cell count base on virus phenotype (Fig. 3D). Among the 24 phenotype-confirmed participants, those with X4-using phenotype had significantly lower CD4 counts compared to those with R5-only phenotype in cluster 4 (141 vs. 440, *p* = 0.003) and in cluster 5 (141 vs. 441, *p* = 0.01) (Fig. 3A and 3D). With well-matched phenotype data, we confirmed that X4-using phenotype is associated with significantly lower CD4 count in CRF01_AE. The higher prevalence of X4 phenotype in cluster 4 during the first year of infection and its association with lower CD4 count explained the reason why cluster 4 had lower CD4 count compared to cluster 5 starting from early infection stage (Fig. 1C-D).

### Genetic and structural determinants for higher CXCR4 usage propensity

To explore the mechanism for higher X4-using propensity exhibited by cluster 4 viruses, we first compared the V3 loop sequences between clusters 4 and 5. We found that cluster 4 viruses have two highly conserved basic amino acids at positions 13 and 32 in its V3 loop (R13 and K32, HXB2 numbering R308 and K327), which were present in only about 8% of the cluster 5 viruses (Fig. 4A-B). These two conserved amino acids confer cluster 4 viruses a higher positively charged V3 loop, upon which fewer mutations may be required to switch from the R5 to X4 phenotype. Structure analysis using V3-docking models (18) suggested that residue R13 in cluster 4 potentially forms salt bridges with D262 and E277 and a hydrogen bond with the side chain of H281 in the CXCR4 coreceptor, while the corresponding residue in cluster 5 may form hydrogen bonds with K22 and D276 in the CCR5 coreceptor (Fig. 4C). Because the ligand binding pocket of CXCR4 is more negatively charged than that in CCR5, the positively charged residue R13 in cluster 4 viruses is more favored. The residue K32 in cluster 4 may form salt bridges with CXCR4’s N terminus, which contains more acidic residues than CCR5’s N terminus (Fig. 4C). Therefore, the highly conserved residues R13 and K32 in cluster 4 may be key determinants for the higher tendency of X4 usage. Interestingly, the V3 region of cluster 4 has less variations compared to the CRF01_AE ancestor sequences from Thailand, notably the preservation of residue K32, while cluster 5 is more divergent (Fig. 4A and SI Appendix, Fig. S5).

**Figure 4.**
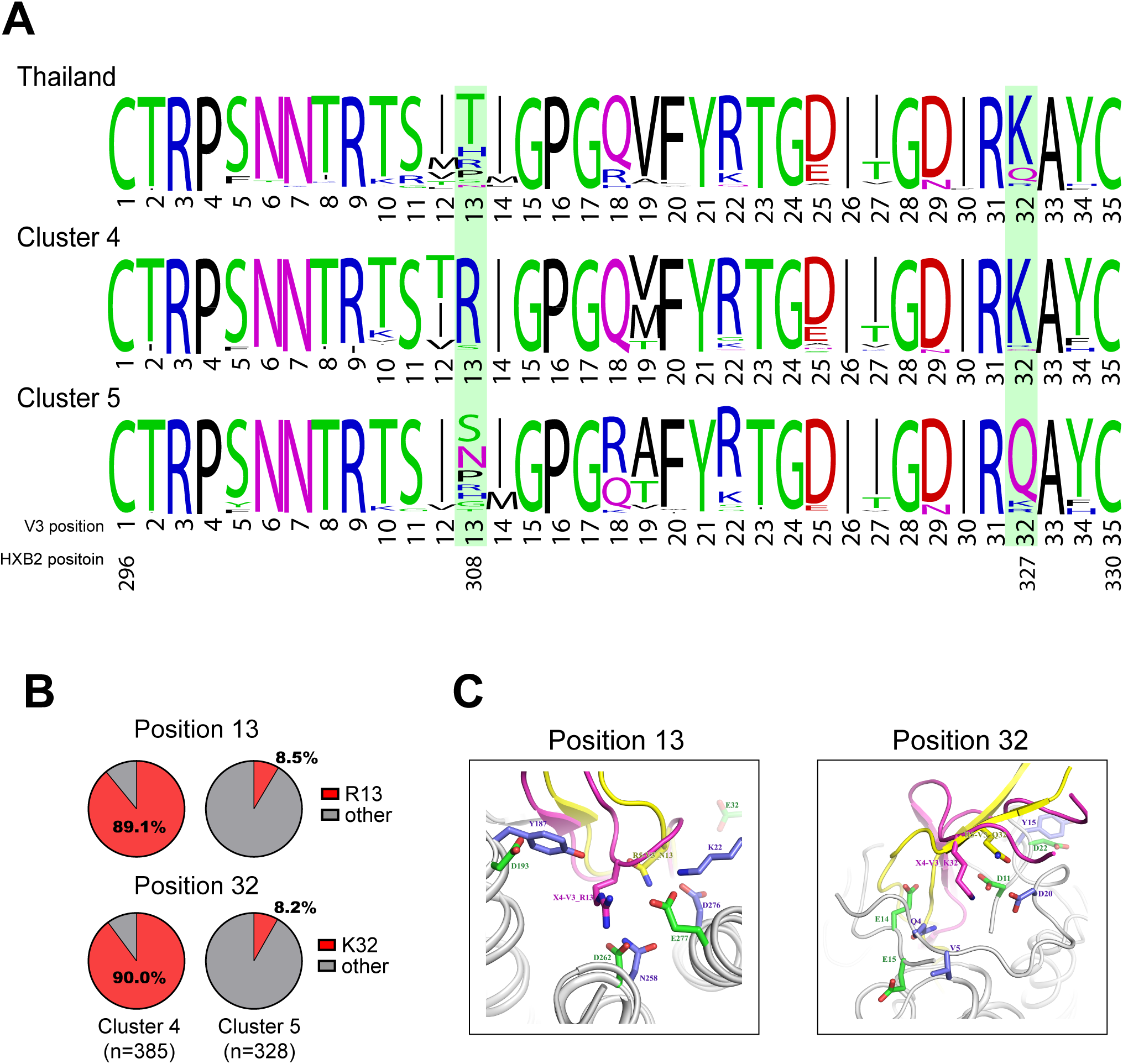
Genetic determinants and structural basis of the higher X4-using tendency in CRF01_AE cluster 4. (A) The frequency of each V3 amino acid was determined with a total of 385 available sequences from CRF01_AE cluster 4, 328 available sequences from CRF01_AE cluster 5, and 34 sequences from Thailand downloaded from the Los Alamos HIV sequence database (before the year 2000). The plots were generated using the WebLogo tool (https://weblogo.berkeley.edu/). (B) Pie charts showing the prevalence of positively charged amino acid K13 and R32 in CRF01_AE cluster 4 and cluster 5. (C) Structural analysis for V3 position 13 and 32 in binding of the CCR5 and CXCR4 coreceptors using the V3-docking model.

Despite the fact that R13 and K32 were present in vast majority of cluster 4 viruses, only about 30% of individuals in cluster 4 were phenotypically confirmed to harbor X4 viruses. This indicates the existence of additional determinants to shift to the X4 phenotype. To further identify key amino acids governing the phenotype switch, we analyzed the genetic composition of PBMC isolates at the single-genome level (Fig. 5 and SI Appendix, Fig. S6 and Table S3). In order to "sieve out” the X4-using variant(s) from the entire viral population (that is, the exact sequence(s) accounting for the X4-using phenotype), we further sequenced the viruses released from the GHOST.X4 cell culture by SGA for the five X4-using samples (SI Appendix, Table S3). Comparing the genetic composition between the PBMC viral isolates and the concurrent plasma viral population showed two different patterns: in majority of R5 subjects (15 of 19), the V3 lineages in the PBMC isolate and in plasma were in proportion, that is, the predominant lineage in the PBMC isolate was also the predominant one in plasma (Fig. 5A and SI Appendix, Table S3). In contrast, in four of the five X4 isolates (with the exception of CYM194), the predominant, phenotypically confirmed X4 lineage in the PBMC isolate existed as minor variant in plasma (Fig. 5B and SI Appendix, Table S3). All V3 lineages detected in the viral isolates from PBMC were present in plasma, and more V3 lineages were detected in plasma than in the primary isolates (Fig. 5 and SI Appendix, Table S3).

**Figure 5.**
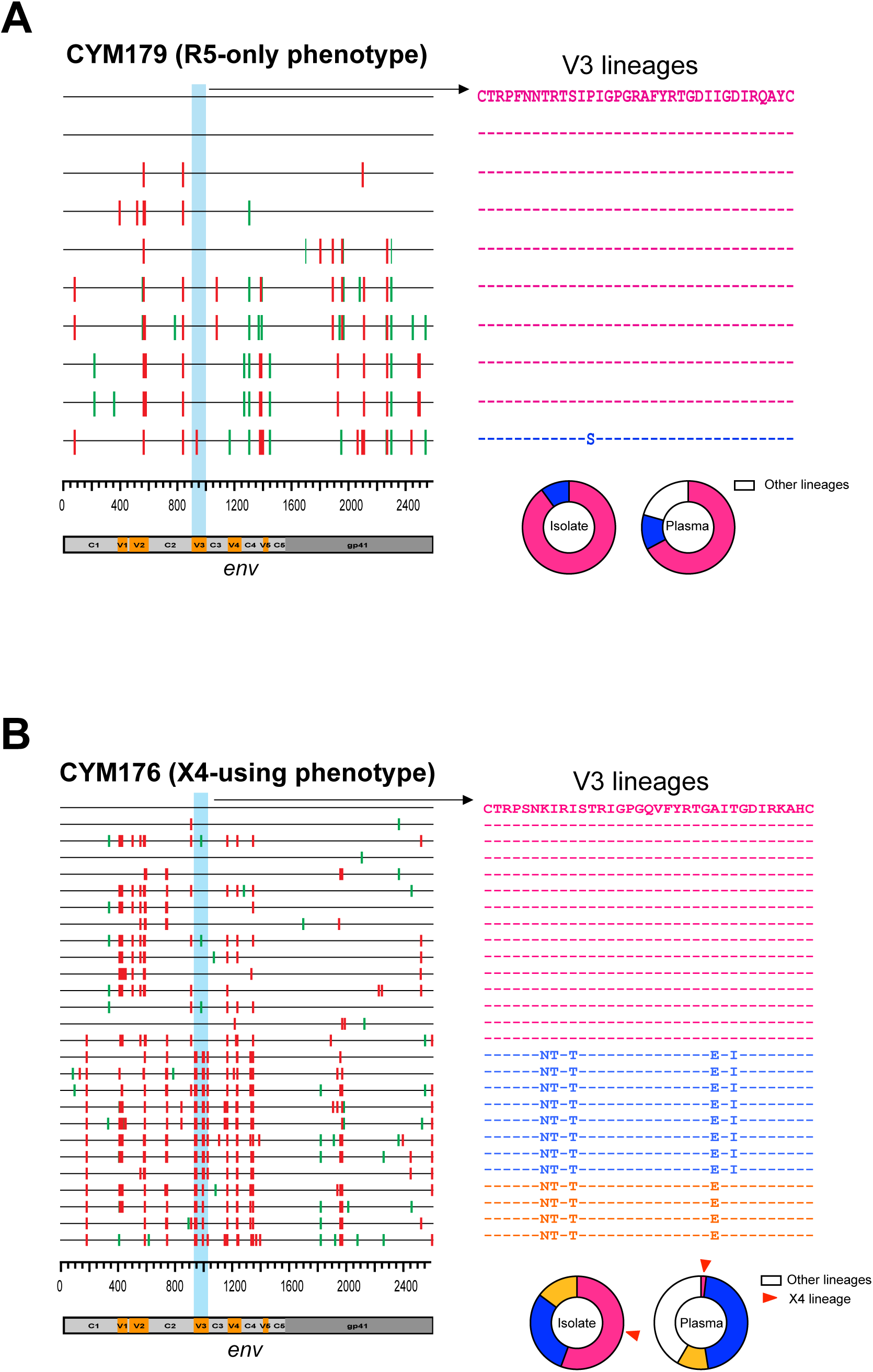
Genetic composition of the PBMC viral isolates. (A-B) SGA-derived gp160 sequences from the R5 isolate CYM179 (A) and the dual tropic isolate CYM176 (B) were shown using highlighter plot. In each sample, one sequence with the predominant V3 lineage was used as the master sequence. The synonymous and non-synonymous substitutions compared to the master sequence were shown in green and red, respectively. The corresponding V3 alignment was shown on the right, and different V3 lineages were color-coded. The circle plots show the proportion of each V3 lineage in the PBMC viral isolate as detected by SGA and in the plasma as detected by deep sequencing. In CYM176, the red arrows indicate the phenotypically confirmed X4 lineage.

Several V3 alterations were found in the phenotypically confirmed X4 sequences. First, all X4-using sequences lost the N-linked glycan site at the beginning of the V3 loop (V3 positions 6–8, HXB2 numbering 301–303), mostly by T to I substitution at position 8 (Fig. 6A). Second, while residue N was invariably found in all R5 sequences at position 7, all but one X4-using sequences had residue K at this position (Fig. 6A). Third, either E or D were found at position 25 in R5 sequences, however, non-E/D substitutions (S/A/G) were present in majority of X4-using sequences (Fig. 6A). Interestingly, none of the X4 sequences have positively charged amino acid R or K at V3 position 11 or 25, which are important for X4 usage in other HIV-1 subtypes (19–21). This implies different evolutionary pathway of coreceptor switching in CRF01_AE HIV-1. In genotypic prediction, all X4-using sequences had Geno2pheno FPR values below 2%, and V3 net charge no less than 5. In contrast, all R5 sequences had FPR values higher than 2%, and V3 net charge no more than 5. As expected, cluster 4 has an overall higher V3 net charge than cluster 5 (Fig. 6A). Notably, five sequences in cluster 4 with FPR below 5% were in fact R5-only phenotype (Fig. 6A). Therefore, using FPR 5% as the cutoff may significantly overestimate the prevalence of X4 phenotype in CRF01_AE.

**Figure 6.**
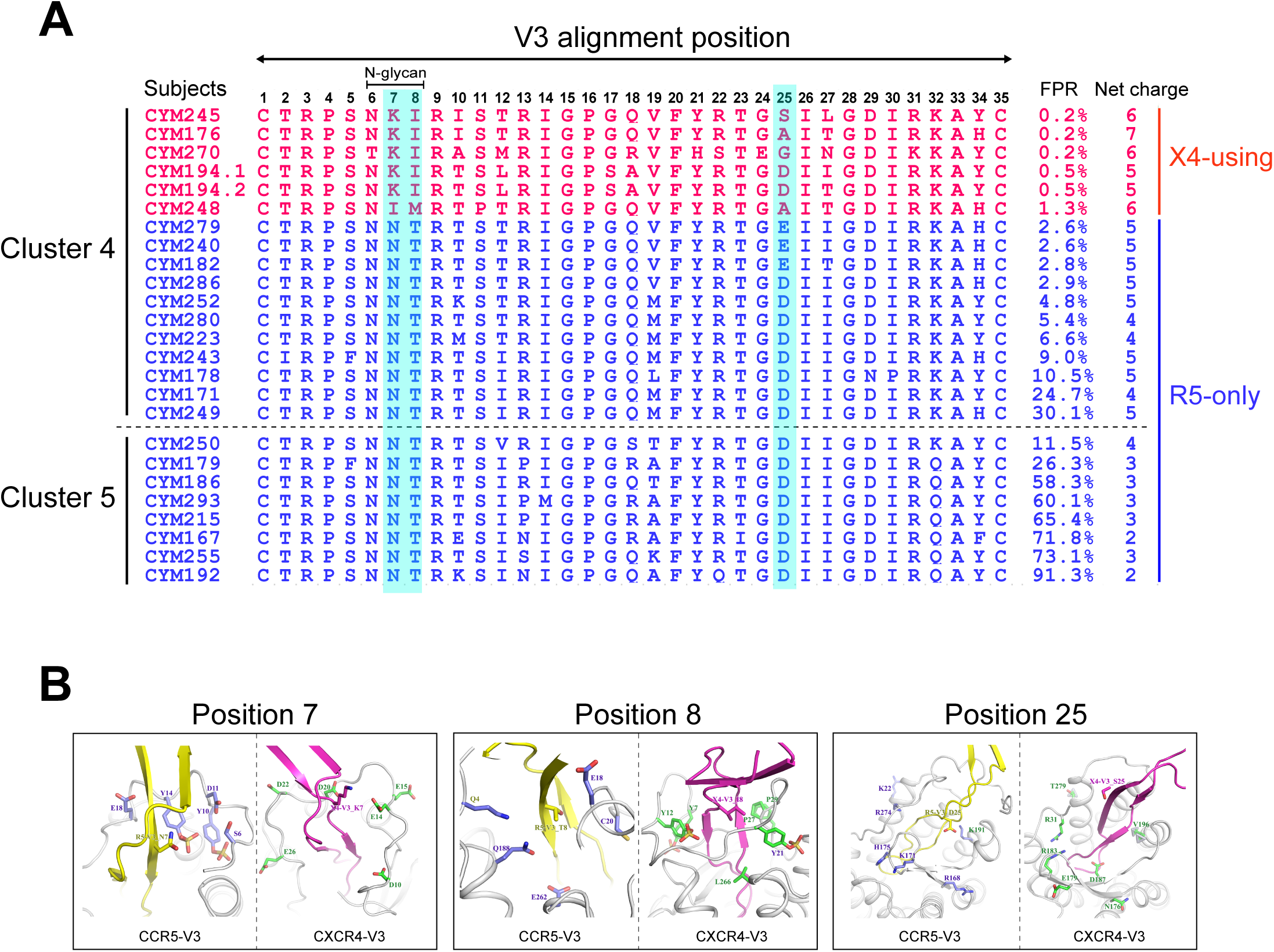
Genetic characteristics of phenotypically confirmed X4 sequences and structural modeling for coreceptor binding. (A) V3 amino acids alignment of SGA-derived sequences from the phenotype-confirmed primary viral isolates. Sequences shown in red were phenotypically confirmed X4-using sequences (that is, sequences sieved out from the GHOST. CXCR4 culture by SGA). Sequences shown in blue were the predominant V3 forms in the R5 isolates. Key V3 positions associated with X4-using phenotype were shaded in light blue. (B) Structural modeling for V3 positions 7, 8 and 25 in binding of coreceptors CCR5 and CXCR4 using the V3-docking model.

Using the CCR5-V3 and CXCR4-V3 complex models (22, 23), we also investigated the role of V3 positions 7, 8 and 25 in viral tropism from a structural perspective. In the model of CCR5-V3 complex, residue T8 in R5 V3 loop is surrounded by hydrophilic amino acids, suggesting that residue T8 is more favored by hydrophilic environment. However, in the model of CXCR4-V3 complex, residue I8 in the X4 V3 loop is surrounded by hydrophobic amino acids (Fig. 6B). This could explain why all X4 viruses have T to I/M substitutions at position 8 because the residue T8 may not well fit the hydrophobic environment within CXCR4’s ligand binding pocket. In the CXCR4-V3 complex, V3 position 7 is surrounded by negatively charged residues, which favor the interaction with positively charged residues K7 in X4 sequences (Fig. 6B). Compared to the corresponding region in the ligand binding pocket of CXCR4, the ligand binding pocket in CCR5 around V3 position 25 contains more positively charged residues, which are favored for interacting with the negatively charged residues D/E25. This explained why all R5 viruses have D/E at V3 position 25. However, D/E may be less favored in the less positively charged environment in the ligand binding pocket of CXCR4 (Fig. 6B). Therefore, non-D/E substitutions as observed in X4-using sequences would be required for efficient X4 binding (Fig. 6B). Taken together, genetic analysis in combination with structural modeling showed that specific V3 substitutions at position 7, 8 and 25 may be required to achieve X4-using phenotype in the context of CRF01_AE cluster 4 envelope.

## Discussion

Since the initial introduction to human in the early 20th century, HIV-1 evolved genetically and biologically with faster pace than other viruses, due to both the high error-prone nature of its reverse transcriptase and the unusual transmission routes, such as drug injections, heterosexual transmission and MSM activities. The Chinese HIV-1 epidemic with multiple subtypes and their genetic clusters circulating at the same time provides a unique opportunity to monitor virus evolution at both the genotype and phenotype levels.

Past studies on HIV-1 evolution were mainly focused on virus genotype not phenotype, because the later takes longer observation time and requires large well-matched samples. In this study, we focused on various genetic clusters of CRF01_AE HIV-1 in China. In the large cross sectional data from the NHMES, we discovered significant difference in CD4 count between people infected by CRF01_AE cluster 4 and 5, at both early and later stage of infection. The lower CD4 count in cluster 4 is directly associated with the higher prevalence of X4 virus based on genotypic prediction. This genotype-based observation in large population was further confirmed by well-matched genotyping and phenotyping data from a MSM sero-incidence cohort. We observed that among sero-convertors, those harboring X4-using viruses had rapid CD4 loss.

It is usually considered that X4 variants emerge during late stage of infection, with the development of immunodeficiency of the host (24). In subtype B HIV-1, around 50% of patients underwent coreceptor switch, usually after 5 years of infection, which correlated with rapid CD4 decline and faster progression to AIDS (16, 25–30). The exact time of coreceptor switch in different HIV-1 subtypes are not well understood. A recent study did not detect X4 variants in subtype B infected people who were within 2 years of infection (31). We demonstrated here that in certain HIV-1 subtypes or clusters, such as CRF01_AE cluster 4, coreceptor switch can occur much earlier than previously thought. This unusually fast speed of coreceptor switch is associated with the rapid CD4 loss in CRF01_AE cluster 4 compared to cluster 5 as well as other HIV-1 subtypes. Supported by genetic and structural analysis, the unique V3 signatures in cluster 4 (R13 and K32), which confer higher V3 net charge may be the major intrinsic determinant for such a high coreceptor switch tendency. Further efforts are required to understand whether the rapid emergence of X4 virus in vivo is essentially a random event due to accumulation of mutations, or also driven by host factor(s) like immune pressure. In addition, envelope positions outside of the V3 loop like V1V2 could also play a part in this process. A better understanding of the driving force and evolutionary pathway for coreceptor switch in CRF01_AE cluster 4 may lead to strategies to block the early emergence of X4 virus during infection.

The genetic features of the X4 variants in CRF01_AE cluster 4 are also different from previously found in other HIV-1 subtypes. In particular, none of the X4 sequences in CRF01_AE cluster 4 have positively charged amino acid (R or K) at V3 positions 11 or 25, which are key amino acids for X4 usage observed in other subtypes (19–21). Instead, all of them lost the V3 glycan (the N301 glycan), and nearly all have residue K at V3 position 7. This highlights different evolutionary pathways for coreceptor switching in different HIV-1 subtypes. N301 glycan has previously been shown to be functionally critical for both coreceptor utilization and virus replication in subtype B (32, 33). With compensatory mutations or a high V3 net charge, loss of the N301 glycan leads to switch from R5 to X4 phenotype (32, 33). However, in the absence of compensatory mutations, loss of this glycan can abolish virus replication (32). Possibility due to this high fitness constraint, N301 glycan is highly conserved in naturally occurring subtype B sequences. A recent study found that in the Los Alamos HIV database, N301 glycan is present in as high as 99% of subtype B sequences with R5 phenotype and more than 80% of sequences with X4 phenotype. Differently, in CRF01_AE, 94% of R5 sequences and 39% of X4 sequences have the N301 glycan site (34). This again indicates that loss of the N301 glycan is an important pathway for coreceptor switch in CRF01_AE HIV-1, but is not a primary route among subtype B viruses. It will be interesting to study in the future whether N301 glycan has a lower fitness barrier in the context of CRF01_AE cluster 4 envelope than in CRF01_AE cluster 5 and other HIV-1 subtypes.

Another interesting finding is the outgrowth of highly replication competent X4 variants in primary viral isolates from CRF01_AE cluster 4. The low frequency of those X4 variants in plasma may not be explained by their low replication fitness, as they rapidly outcompete other lineages in the in vitro setting. Instead, it is more likely due to the compartmentalization of the R5 and X4 viruses in different cell subsets or tissues in vivo. Due to the differential expressions of CCR5 and CXCR4 coreceptors in memory and naïve CD4 T cell subsets (35, 36), R5 and X4 viruses are considered to preferentially replicate in the memory and naïve CD4 subsets, respectively (37–41). It has been shown that naïve CD4 T cells produce viruses at a lower propagation rate than memory T cells (42, 43), possibility due to the relatively low division rate (44, 45). Therefore, in vivo, those minor X4 lineages might compartmentalize in cell subsets or tissues that shut the viruses less efficiently into the blood. Regardless of the mechanism, this observation has its clinical implication: because conventional sequencing method may not be sensitive enough to capture those minor X4 variants in plasma, deep sequencing or phenotypic assay would be required to determine the existence of X4 variants in vivo, especially when using treatment regimens including the CCR5 inhibitor.

In summary, we for the first time demonstrated that various phylogenetic clusters of the same HIV-1 subtype can have disparate pathogenicity and cause different disease outcomes, which filled the missing link between HIV-1 phylogenetic cluster and viral phenotype. At the phenotype level, CRF01_AE cluster 4 evolved enhanced X4 tropism and viral pathogenesis, while cluster 5 became more attenuated due to a decreased potential of using CXCR4. Whether the process of "phenotype divergence” occurred as a random founder event from the initial seeding clusters, or due to adaptation to different hosts or transmission routs remains to be studied. Our study emphasizes the importance of monitoring HIV-1 genetic drift and phenotype shift at the phylogenetic cluster level in order to timely control the spread of more pathogenic viruses like CRF01_AE cluster 4.

## Materials and Methods

### Study participants

The study participants were from the national HIV molecular epidemiology survey (NHMES) and the Beijing Chaoyang District MSM cohort (the CYM cohort). Written informed consent was obtained from all study participants. See SI Appendix, Supplementary Text for details.

### Enzyme Immunoassay (EIA)

In order to distinguish recent HIV-1 infections from long-term HIV-1 infections, the Enzyme Immunoassay (EIA) was performed using the Maxim HIV-1 Limiting Antigen-Avidity (LAg-Avidity) EIA kit (Maxim Biomedical). The experiment and data analysis were performed according to manufacturer’s instructions (Maxim Biomedical).

### Viral RNA extraction and cDNA synthesis

Viral RNA was extracted from 200 μl of plasma sample using the QIAamp Viral RNA Mini Kit (Qiagen). RNA was eluted into 50 μl of RNase free water. A total of 17 μl viral RNA was used for cDNA synthesis using the SuperScript III reverse transcriptase (Invitrogen) with the Oligo (dT) primer. The cDNA was immediately used for PCR amplification.

### Library preparation for sequencing on Illumina MiSeq

The Illumina MiSeq library was prepared using a nested PCR approach. The 8 nt Illumina index and adaptors (P5 and P7) were added to both ends of the second round PCR primers (SI Appendix, Fig. S2 and Table S4). The second round PCR products were gel-purified to remove unspecific bands and primer dimers, and quantified by qPCR using the KAPA SYBR FAST qPCR kit according to manufacturer’s instructions (KAPA Biosystems). See SI Appendix, Supplementary Text for details.

### Next generation sequencing and data analysis

The pooled DNA library was sequenced on an Illumina MiSeq using the MiSeq Reagent Kit v2 (500 cycles, Illumina) as previously described (46). Each pair of fastq reads in files "read 1” and "read 2” were merged by the FLASH software (47). The merged fastq files were then filtered based on data quality on the Galaxy server (48) using the following parameter: no more than 10 bases with Q score lower than 30 in each read. The filtered clean reads were then converted into fasta format. In each individual, identical reads were collapsed into haplotypes after the primer regions were trimmed. The frequency of each haplotype among the total clean reads was calculated. The most frequent haplotype in each individual was used to infer the phylogenetic relationship among all deep sequenced individuals.

### Genotypic prediction of co-receptor usage

Genotypic prediction of co-receptor usage was performed using the Geno2pheno clonal model (https://coreceptor.geno2pheno.org) (49). For the deep sequencing data, the FPR (false positive rate, the probability of classifying an R5-virus falsely as X4) was obtained for each V3 haplotype. To avoid the impact of potential PCR or sequencing error on data analysis, only V3 haplotypes appeared three times or more in each sample were used for analysis, while V3 singletons and those appeared only twice where discarded. The frequency distribution of FPR value in each sample was obtained by calculating the frequency of each V3 haplotype among the total reads analyzed.

### Primary virus isolation from PBMC

To obtain primary virus isolates, cryopreserved PBMC from HIV-1 infected patients were co-cultivated with stimulated normal PBMC from healthy donors. In brief, fresh PBMC from healthy donors were stimulated for 3 days in RPMI1640 containing 10% fetal bovine serum (FBS), interleukin 2 (IL-2) (32 U/ml; PeproTech), soluble anti-CD3 (0.2 μg/ml; eBioscience) and soluble anti-CD28 (0.2 μg/ml; eBioscience) as described previously(50). After stimulation, cells were washed twice with RPMI1640 to remove the residue simulating antibodies. A total of 10^7^ stimulated PBMC from healthy donor were then mixed with 10^7^ of PBMC from an infected patient. The cell mixtures were then depleted for CD8 T cells using the EasySep Human CD8 Positive Selection Kit (Stemcell Technologies). The CD8-depleted cell mixture was cultured in a T25 flask with RPMI1640 containing 10% fetal bovine serum (FBS) and 32 U/ml IL-2 (PeproTech) for up to 4 weeks. Every 3 days, half volume of the culture supernatant was replaced with fresh medium. Every 7 days, half of the entire culture (including cells) was removed, and 5×10^6^ of stimulated, CD8 T cell depleted PBMC from healthy donors were added. The p24 concentration in the culture supernatant was measured every week. Majority of cultures achieved peak p24 production around week 3. Cultures with p24 concentration less than 2 ng/ml at week 4 were considered to be failed.

### Coreceptor tropism determination

Coreceptor tropism of the primary viral isolates was determined using the GHOST(3).CCR5 and GHOST(3).CXCR4 cell lines (51). Both GFP expression in the Ghost cell lines and viral p24 production were used to determine the coreceptor usage. See SI Appendix, Supplementary Text for details.

### Single genome amplification

Single genome amplification (SGA) was performed as previously described (52). The sequences were aligned using GeneCutter (https://www.hiv.lanl.gov/content/sequence/GENE_CUTTER/cutter.html), followed by the manual adjustment to obtain the optimal alignment. See SI Appendix, Supplementary Text for details.

### Statistical analysis

All statistical analysis was performed using the Prism 7 (GraphPad Software). Statistical differences were determined using two-tailed Mann-Whitney test or Fisher’s exact test as indicated in the figure legends. The exact *p* values were provided in the figures.

### Data availability

Newly generated nucleic acid sequences in the current study were deposited in GenBank with accession numbers MH672692-MH673032.

## Acknowledgements

We thank Dr. Cecilia Cheng-Mayer for comments on the manuscript. This work was supported by China National Major Project for Infectious Diseases Control and Prevention, and China Key Project of the State Key Laboratory of Infectious Diseases Control and Prevention.

## Author contributions

YS conceived and designed the study. HS, WO, YF, JZ, FL, JH and HP performed experiments. QT and BW designed and performed the structural analysis. HS, WO, YF, HX, LM, QT, BW and YS analyzed the data. HS, BW and YS wrote and edited the manuscript.

